# Induced pluripotent stem cell model revealed impaired neurovascular interaction in genetic small vessel disease CADASIL

**DOI:** 10.1101/2023.04.26.538393

**Authors:** Wenjun Zhang, Xiangjun Zhao, Xuewei Qi, Susan J Kimber, Nigel Hooper, Tao Wang

## Abstract

Cerebral Autosomal Dominant Arteriopathy with Subcortical Infarcts and Leukoencephalopathy (CADASIL) is the most common genetic small vessel disease caused by variants in the *NOTCH3* gene. Patients with CADASIL experience recurrent strokes, developing into cognitive defect and vascular dementia. CADASIL is a late-onset vascular condition, but migraine and brain MRI lesions appear in CADASIL patients as early as their teens and twenties, suggesting an abnormal neurovascular interaction at the neurovascular unit (NVU) where microvessels meet the brain parenchyma. To understand the molecular mechanisms of CADASIL, we established induced pluripotent stem cell (iPSC) models from CADASIL patients and differentiated the iPSCs into the major NVU cell types including brain microvascular endothelial-like cells (BMECs), vascular mural cells (MCs), astrocytes and cortical projection neurons. We then built an *in vitro* NVU model by co-culturing different neurovascular cell types in Transwells and evaluated the blood brain barrier (BBB) function by measuring transendothelial electrical resistance (TEER). Results showed that, while the wild-type MCs, astrocytes and neurons could all independently and significantly enhance TEER values of the iPSC-BMECs, such capability of MCs from iPSCs of CADASIL patients was significantly impeded. Additionally, the barrier function of the BMECs from CADASIL iPSCs was significantly impaired, accompanied with disorganised tight junctions in iPSC-BMECs, which could not be effectively rescued by the wild-type MCs, astrocytes and neurons. Our findings provide new insight into early disease pathologies on the neurovascular interaction and BBB function at the molecular and cellular levels for CADASIL, which helps inform future therapeutic development.

## 1. INTRODUCTION

CADASIL (Cerebral Autosomal Dominant Arteriopathy with Subcortical Infarcts and Leukoencephalopathy) is a genetic small vessel disease (SVD) caused by heterozygous variants in the *NOTCH3* gene (Joutel et al., 1996, Sharma et al., 2001). Although a rare condition, CADASIL represents the most common type of monogenic SVD with a prevalence of 2 to 5 per 100,000 people and a likelihood of being underdiagnosed (Rutten et al., 2016, Narayan et al., 2012, Razvi et al., 2005). Patients with CADASIL usually experience recurrent ischemic strokes, migraines with aura, psychiatric disturbances and seizures, and gradually develop to cognitive impairment and vascular dementia (Joutel et al., 1997, Chabriat et al., 2009, Adib-Samii et al., 2010, Charlton et al., 2006). Despite excellent research carried out in this field, molecular mechanisms underlying this condition are still not entirely clear, therefore, no specific and effective treatments are available to date.

Typical pathological changes observed in CADASIL patients include vascular smooth muscle cell (VSMC) degeneration in small arteries, accumulation of the extracellular domain of NOTCH3 protein, deposition of granular osmiophilic materials (GOM) around VSMCs, and thickening of small blood vessel walls (Joutel et al., 2001, Joutel et al., 1997, Chabriat et al., 2020). However, the mechanisms by which vascular dysfunction leads to clinical manifestations in the brain has not been fully understood. Recurrent strokes in CADASIL patients would certainly damage brain function, contributing to the development of cognitive defects and the eventual dementia. However, CADASIL is a late-onset condition where strokes usually occur between 40 and 50 years of age. According to the natural history of CADASIL patients, migraine with aura and abnormalities of white matter on T2 weighted MRI images happen as early as in their teens and twenties, well before the onset of strokes (Chabriat et al., 2009, Desmond et al., 1999). This suggests a likelihood of defects at the interface where blood vessels meet the brain parenchyma.

In the central nervous system (CNS), functional connections between blood vessels and neurons rely on the unique structure – the neurovascular unit (NVU) comprising mainly brain microvascular endothelial cells (BMECs), mural cells (MCs, including pericytes on capillaries and VSMCs on arterioles), astrocytes, and neurons. BMECs are surrounded by MCs that are embedded in the basement membrane, and the microvessels are ensheathed by astrocyte endfeet that bridge to the neurons and are surrounded by microglia (Andreone et al., 2015). Complex and dynamic interactions between these neurovascular cell types regulate the blood-brain barrier (BBB) function and cerebral blood flow (CBF), which helps to maintain the homeostasis of the CNS (Abbott et al., 2006, Daneman and Prat, 2015). Therefore, the NVU, as the neuro-vascular interface, is a key target for understanding the molecular mechanisms of CADASIL and SVD in general, which could potentially inform future drug development.

The BMECs are the central component of the BBB structure and have unique properties distinct from ECs in peripheral vasculatures. BMECs have a low rate of transcytosis, a lack of fenestrations, and are particularly enriched in tight junction proteins (e.g., claudin-5, occludin, ZO-1) that are essential in reducing paracellular diffusions and contribute to the barrier function of BBB (Daneman and Prat, 2015, Coomber and Stewart, 1985, Wallez and Huber, 2008). BMECs have high numbers of transporters. The glucose transporter (GLUT-1) is critically important for brain function by maintaining the continuous glucose level and energy demands to support neural cell growth and activities (Yazdani et al., 2019). The ATP-binding cassette (ABC) efflux transporter family members (e.g., p-glycoprotein) are responsible for eliminating harmful substances from the brain into the blood (Chai et al., 2022). The highly polarised nature of BMECs ensures efficient transportation to occur between the brain and blood. The expression of leukocyte adhesion molecule is also low in BMECs, which greatly limits the entry of peripheral immune cells into the CNS, thus protecting the brain from damage when inflammation occurs (Daneman and Prat, 2015, Daneman et al., 2010a). The tight barrier property of BMECs creates a high trans-endothelial electrical resistance (TEER) of >1000 Ω.cm^2^, which is commonly measured using a Voltmeter in experimental settings *in vitro* (Newman et al., 2020, Neal et al., 2019). The BBB permeability can also be determined by observing the diffusion of fluorescent small molecules across the BMEC monolayer (Stebbins et al., 2016).

The unique barrier property of BMECs is not intrinsic to the cells but induced by the CNS environment (Janzer and Raff, 1987). Other neurovascular cell types in the NVU significantly strengthen the barrier function of BMECs (Hongjin et al., 2020, Daneman and Prat, 2015). Neural progenitor cells (NPCs) regulate CNS vascularisation in the developing brain (Tata and Ruhrberg, 2018), and the neuronal activity is also important in the establishment of the efflux transporters of the BBB (Pulido et al., 2020). Astrocytes enhance the BBB barrier function through their perivascular endfeet and the secretion of a range of soluble factors (Alvarez et al., 2013). Capillaries in the CNS have significantly higher pericyte coverage than the peripheral vascular bed, which contributes to the regulation of the BBB (Brown et al., 2019). However, the functional interactions between the neurovascular cell types have not been addressed in CADASIL at the cellular level. We hypothesised that impairments of the neurovascular interaction at the NVU contribute to the development of the CADASIL phenotype.

We have previously established induced pluripotent stem cell (iPSC) models from CADASIL patients and demonstrated an intrinsic defect of MCs in supporting angiogenic tubule structures, likely via a downregulation of PDGFRβ in MCs and a detrimental effect to the adjacent ECs (Kelleher et al., 2019). This iPSC model was used in this study to uncover the neurovascular interaction in CADASIL. We differentiated the iPSCs into BMEC-like cells (BMECs), astrocytes and cortical projection neurons, in addition to VSMCs and ECs, and demonstrated that iPSC-derived MCs, astrocytes and neurons could all significantly enhance the TEER values of BMECs, but such capability was significantly blunted for the CADASIL iPSC-derived MCs. The BBB barrier function was also significantly damaged in the CADASIL iPSC-derived BMECs, which could not be fully rescued by the wild-type iPSC-MCs, astrocytes, or neurons.

## 2. MATERIALS AND METHODS

### 2.1 Cell lines and cell culture

The CADASIL iPSC lines were established from skin biopsies of two CADASIL patients carrying *NOTCH3* variants Arg153C and Cys224Try, respectively, as reported in our previous study (Kelleher et al., 2019). The five control iPSC clones were from three healthy individuals with two iPSC clones (02C3 and 02C9) reported by Kelleher (Kelleher et al., 2019), two clones (SW171a and SW174a) reported by Wood (Wood et al., 2020, Woods et al., 2020), and OX1-19 line reported by Jarosz-Griffiths (Jarosz-Griffiths et al., 2019). Prior to differentiation, all iPSCs were cultured in Vitronectin Recombinant Human Protein (VTN-N) (ThermoFisher, A14700) pre-coated 6-well plates with 2 mL TeSR-E8 medium (Life Technologies, 05990) at 37°C with 5% CO2, except for the OX1-19 iPSCs that were cultured on Matrigel (Corning, 354277) pre-coated 6-well plates in mTeSR1 complete medium (85850, STEMCELL Technologies).

Primary human coronary arterial endothelial cells (HCAECs) (PromoCell, C-12222) were cultured in 6-well plates (Corning, 3516) with 2 mL Endothelial Cell Growth Medium (MV2, PromoCell, C-22121) and maintained at 37°C incubator with 5% CO2.

### 2.2 Differentiation of iPSCs into brain microvessel endothelial-like cells (BMECs)

IPSCs were seeded on Matrigel pre-coated 6-well plate at a cell density 25,000-400,000 per well in TESR-E8 medium, termed day 0. From day 0 to day 5 the medium was changed daily with 2 mL DMEM/F-12 containing 20% KnockOut™ Serum (ThermoFisher), 1 x Non Essential Amino Acid (NEAA, Life Technologies, 11140), 0.5 x L-glutamine and 0.1 mM β-mercaptoethanol. On day 6, the medium was replaced by Human Endothelial serum free medium (hESFM, Invitrogen, 11111) containing 1% Human Platelet Poor Plasma-derived Serum (SH, Sigma, P2918), 20 ng/mL FGF2 and 10 uM All-trans Retinoic Acid (RA, Sigma, R2625) and cultured for two days. On day 8, BMECs were sub-cultured onto plates or Transwells that were pre-coated with solution of Collagen IV (Sigma, C5533, 1 mg/mL in acetic acid)/fibronectin (Sigma, F1141)/water (C/F/W) to be 4:1:5 for purification of BMECs and measurement of transendothelial electrical resistance (TEER).

### 2.3 Differentiation of iPSCs into cortical projection neurons

IPSCs differentiation into cortical projection neurons was adapted from a protocol by Shi et al (Shi et al., 2012). IPSCs were dissociated with EDTA and seeded on to 12-well plates (Corning) pre-coated with Matrigel and cultured in TESR-E8 medium. When reaching 100% confluency, the culture medium was changed to neural induction medium (NIM) containing 1 μM Dorsomorphin (3093, Tocris) and 10 μM SB431542 in neural maintenance medium (NMM). NMM was a 1:1 mixture of N2 (Thermo Fisher Scientific, 17502001) and B27 medium (Thermo Fisher Scientific, 17504044). N2 medium consisted of DMEM/F-12 GlutaMAX medium (Life Technologies, 31331), 1 x N2, 5 μg/mL insulin (Sigma, I9278), 1 mM L-Glutamine (Life Technologies, 25030024), 100 μM NEAA, 100 μM 2-mercaptoethanol (Life Technologies, 31350) and 0.5% Penicillin-Streptomycin (10,000 U/mL) (Life Technologies, 15140). B27 medium consists of 1 x Neurobasal medium (Life Technologies, 12348), 1 x B27, 200 mM L-Glutamine and and 0.5% Penicillin-Streptomycin. The medium was changed daily.

On day 10-12 after induction, the cells were passaged with 0.5 mL Dispase (Life Technologies, 17105) and transferred onto Laminin (Sigma, L2020) pre-coated 6-well plate and cultured in 2 mL fresh NIM. After 24 hours, medium was changed to NMM containing 20 ng/mL Recombinant Human FGF-basic (FGF2) (PeproTech, 100-18C) for 3-4 days. FGF2 was then withdrawn when neural rosettes appeared, and cells were maintained in NMM. Rosettes were isolated from cultures using Neural Rosette Selection Reagent (Stemcell Technologies, #05832), this was the NPC stage.

On day 25 after induction, cells were dissociated into single cells with Accutase (Stemcell Technologies, #07920) and transfered into a new Laminin pre-coated plate with medium changed every other day. Mature neurons after day 60 were used for all experiments.

### 2.4 Differentiation of iPSCs into astrocytes

During neuron differentiation, selected neural rosettes were seeded onto Laminin pre-coated 6-well plate and cultured with 2 mL of NMM at 37°C for 24 hours, this was marked as day 0 of astrocyte differentiation. On day 1, medium was changed to 2 mL of complete STEMdiffTM Astrocyte Differentiation Medium (ADM) (Stemcell Technologies, #08540). On day7 and day 14, cells were passaged with Accutase and seeded onto Laminin pre-coated plate at a density of 1 × 10^5^ cells / cm^2^ and cultured in ADM that was changed every other day. On day 21, cells were passaged and seeded as above and cultured in complete STEMdiffTM Astrocyte Maturation Medium (AMM) (Stemcell Technologies, #08550) with medium changed every other day. Cells were further passaged on days 28 and 35. After day 35, mature astrocytes were observed.

### 2.5 Differentiation of iPSCs into mural cells (MCs) and endothelial cells (ECs)

The differentiations of MCs and ECs from iPSCs were described in our previous publication (Kelleher et al., 2019). Briefly, iPSCs were seeded on to VTN-N coated plates. For MC differentiation via neuroectoderm, TeSR-E8 medium was replaced with Essential 6 medium (E6, Life Technologies, A1516401) supplemented with 10 ng/mL FGF2 and 10 μM SB431542 (Tocris, 1614). The medium was changed every day until day 8 when medium was replaced by E6 medium supplemented with 10 ng/mL human Platelet-Derived Growth Factor BB (PDGF-BB, PeproTech, 100-14B) and 2 ng/mL Recombinant Human Transforming Growth Factor-β1 (TGF-β1, PeproTech, 100-21). Cells were passaged when reaching 70% confluency until day 18. For EC differentiation, E8 medium was replaced with E6 medium supplemented with 6 uM CHIR99021 (Tocris Bioscience, 4423), 20 ng/mL Bone Morphogenetic Protein 4 (rBMP4, (PeproTech, 120-05) and 10 ng/mL FGF2 and cultured for 3 days. On day 3, medium was replaced with E6 supplemented with 50 ng/mL vascular endothelial growth factor (VEGF, (PeproTech, 100-20), 10 ng/mL FGF2 and 20 ng/mL rBMP4, and the medium was changed on day 5. On day 7, cells were cultured in E6 medium supplemented with 50 ng/mL VEGF, 10ng/mL FGF2 and 10 μM SB431542, and the medium was changed every other day. On day 12, endothelial cells were purified using a CD31 MicroBead Kit (Miltenyi Biotec, 130-091-935) and further cultured in the Promocell endothelial cell medium (MV2) until use.

### 2.6 *In vitro* neurovascular unit (NVU) model and transendothelial electrical resistance (TEER) measurement

On day 8 of the BMEC differentiation as described in 2.2 above, the iPSC-BMECs were dissociated using Accutase (Invitrogen, A1110501), 1.1 × 10^6^/cells were seeded on to the C/F/W pre-coated 0.4 µm pore diameter 12-well Transwell filter and cultured in the Transwell. From day 9, medium was changed to hESFM medium with 1% HS (without FGF2 and RA) and TEER values were recorded using an Epithelial Volt/Ohm Meter EVOM2™ (EVOM with STX2 electrodes, World Precision Instrument) on days 24, 48, 72, 96 and 120 hours after cell seeding. TEER measurements were normalised by subtracting the background (TEER from a blank well) and then multiplied by the surface area (1.12 cm^2^) of the Transwell filter and presented as Ω/cm^2^. All TEER experiments were performed with at least 3 triplicate wells, and at least three independent differentiations.

### 2.7 Angiogenesis tubule formation assay

IPSC-ECs at day 14 or iPSC-BMECs at day 10 of differentiation were used for the tubule formation assay. Ten thousand HCAECs, HUVECs, iPSC-ECs or BMECs in 200 µL cell culture medium were plated onto 96-well plates pre-coated with a thin layer of Matrigel (50 µL per well), respectively. HCAECs were seeded in 200 µL Endothelial Cell Growth Medium MV2. IPSC-ECs were seeded in E6 medium supplemented with 5 ng/mL VEGF-165 and 2 ng/mL FGF2. IPSC-BMECs were seeded in hESFM with 50 ng/mL VEGF-165 without RA or FGF2. Phase-contrast images were acquired at 3, 6, 24, 48 and 72 hours of culture in a CO2 incubator on an EVOS™ XL Core Imaging System microscopy (AMEX1000, ThermoFisher Scientific) using 4x objectives.

### 2.8 Low density lipoprotein uptake assay

Differentiated BMECs at day 10 were analysed with the Dil-ac-LDL dye (J65597, Alfa Aesar). Culture medium was aspirated then incubated in medium containing 12 μg/mL Dil-ac-LDL for 4 h at 37°C, 5% CO2. Cells were then washed three times with PBS and fixed with 4% paraformaldehyde for 10 minutes. Images were taken with a fluorescent microscope with an excitation wavelength of 594 nm.

### 2.9 Calcium wave imaging of astrocytes

At day 100 of the differentiation, the iPSC-derived astrocytes were labelled using 20 μM Ca2+ indicator dye Fluo-4 AM (Invitrogen, ThermoFischer Scientific) for 30-60 minutes depending on absorption rate of individual cells until the basal fluorescence of fluo-4 in all cells plateaued to a constant level. Recordings excitation/emission wavelengths 494/506 nm were made by confocal time-lapse microscopy under a Zeiss LSM 880 Biolumi confocal laser scanning microscope with a 40x oil objective (Carl Zeiss, UK). Spontaneous activity of cells was recorded for 10 minutes. Five random fields were chosen under microscopy and averaged for each sample. Time-lapse recording and images were analysed using Zeiss microscopy software ZEN, MicroExcel and ImageJ.

### 2.10 Immunofluorescent staining

Immunofluorescent staining was performed as previously described (Wang et al., 2007). Briefly, Cells were fixed with 4% paraformaldehyde, permeabilised with 0.2% Triton X-100 for 3-10 minutes at room temperature, and incubated with blocking buffer (10% donkey serum). The cells were then incubated with primary antibodies with appropriate dilutions (**Suppl. Table 1**) in blocking buffer for 1 hour at room temperature or overnight at 4°C, followed by incubating with fluorescent conjugated secondary antibodies (**Suppl. Table 1**) and conterstained by DAPI. Images were collected on an Olympus BX51 upright microscope using an objective (20x / 0.5 UPlan FLN) and captured using a Coolsnap ES2 camera (Photometrics) through Metavue v7.8.4.0 software (Molecular Devices). Images were then processed and analysed using Fiji ImageJ.

### 2.11 Quantitative real time polymerase chain reaction (qRT-PCR)

QPT-PCR was carried out according to previous report (Kelleher et al., 2019). Briefly, Total RNA was extracted from cells using an RNeasy mini kit (Qiagen, 74104) and reverse transcribed into cDNA. QRT-PCR was performed using PowerUp™ SYBR™ Green Master Mix (A25742, Thermofisher) with PCR cycles of 95°C for 10 minutes, followed by 40 cycles of 95°C for 15 seconds and 60°C for 1 minute. *GAPDH* was used as endogenous control. Data was analysed using 2^-ΔΔCt^ or the Pfaffl method where differences in primer efficiencies were accounted. Primers used in the study are listed in **Suppl. Table 2**.

### 2.12 Statistics

Data were presented as mean ± standard error (SEM). GraphPad PRISM was used for data analysis and statistical tests. Comparisons of two groups were made using an unpaired t-test. For comparisons of data sets above two samples, one-way or two-way ANOVA with Tukey’s multiple comparisons post-hoc test was used.

## 3. RESULTS

### 3.1 Neurovascular interactions enhance the barrier function of iPSC-BMECs but mural cells from CADASIL patients fail to do so

BMEC-like cells were differentiated from iPSCs according to published protocols (**Figure 1A**). During differentiation, the iPSCs gradually lost the typical colonised growth nature of stem cells and acquired cobblestone morphology (**Figure 1B**). The iPSC-BMECs displayed endothelial markers (CD31 and VE-cadherin), tight junction proteins (occludin, claudin5 and ZO1), as well as the glucose transporter (Glut1) on immunofluorescent staining (**Figure 1C**). Functional assays revealed that the iPSC-BMECs had the ability to take up low-density lipoproteins (LDL) (**Figure 1D**) and formed vessel network structures when cultured on Matrigel (**Figure 1E**). To determine the BBB function, the trans-endothelial resistance (TEER) value was recorded across the iPSC-BMEC monolayer in a Transwell setting (**Figure 1F**), which showed a peak TEER ∼1,000 Ω.cm^2^. The barrier function of iPSC-BMECs was compared with that of both iPSC-derived peripheral ECs and human coronary artery endothelial cells (HCAECs), which showed that the TEER values of iPSC-BMECs were significantly higher than the ECs (**Figure 1G**).

**Figure 1.**
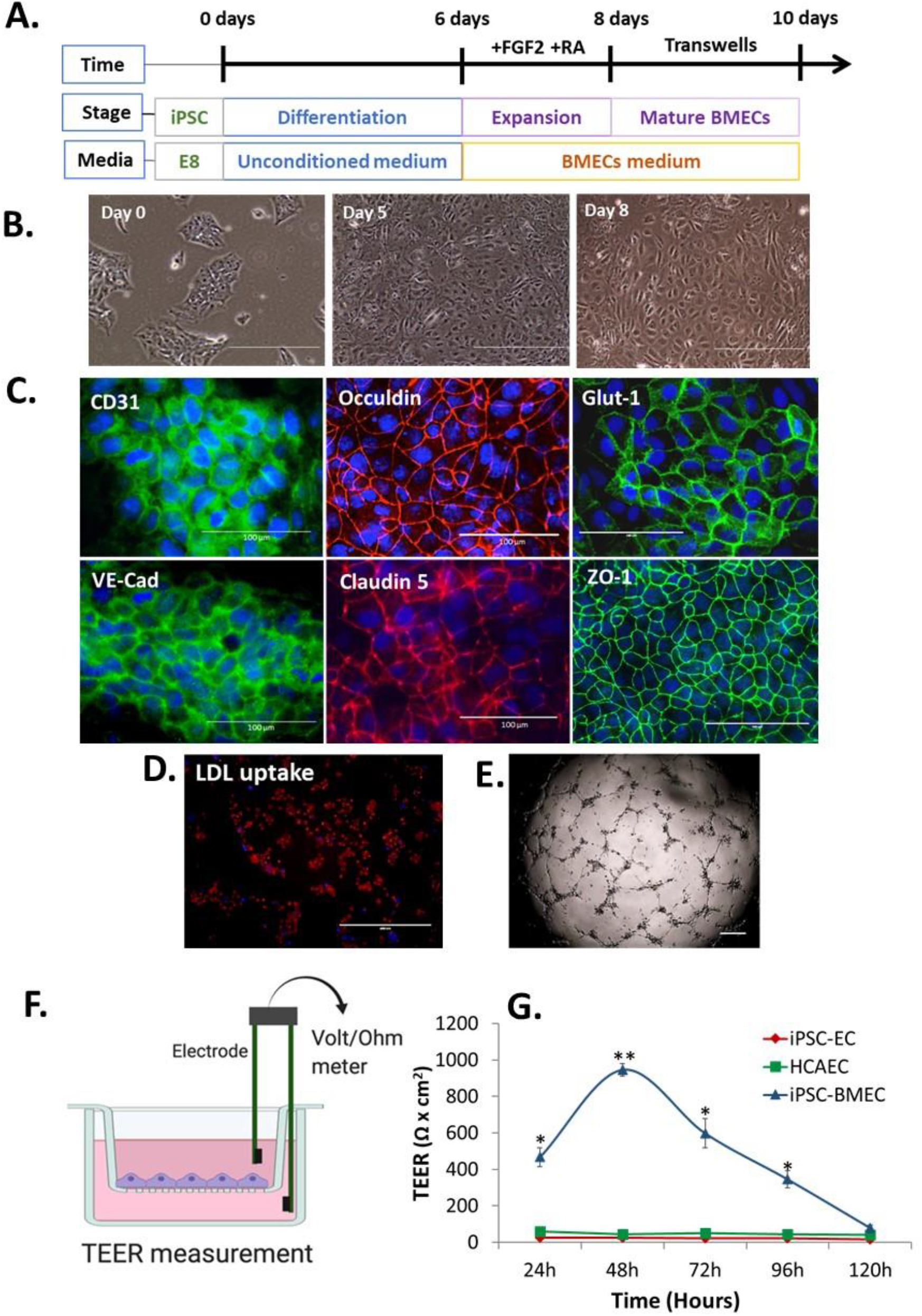
BMEC differentiation from iPSCs and characterisations. **A**. Schematic illustration of BMEC differentiation protocol. **B**. Morphological changes of iPSC during BMEC differentiation under light microscope. **C**. Immunofluorescent staining of BMEC marker proteins, including adherence junction proteins (CD31, VE-cadherine), tight junction proteins (Occludin, claudin 5, ZO-1), and glucose transporter, Glut-1. Scale bars, 100 µm. **D**. IPSC-BMECs at day 10 differentiation were treated with the Dil-ac-LDL dye and visualised under fluorescent microscope. Scale bar, 200 µm. **E**. In vitro angiogenesis assay on Matrigel. Scale bar, 200 µm. **F**. Schematic illustration of the *in vitro* NVU using Transwell setting. The BBB function is measured as TEER values using a Voltmeter. **G**. IPSC-BMECs, iPSC-ECs and hCAECs were seeded on to the insert of the Transwell, respectively. TEER values were measured each day for five consecutive days (120 h). Data are represented as mean ± SEM, n=3. Two-way ANOVA with Tukey’s post hoc test, *p<0.05, **p<0.01. Figure F was created using BioRender.

We then moved on to determine if our iPSC-derived neurovascular cell types could enhance the barrier function of iPSC-BMECs. MCs were differentiated from iPSCs via the neuroectoderm according to our previous report (Kelleher et al., 2019). Differentiation of astrocytes and cortical projection neurons from iPSCs are shown in **Suppl. Figure 1 and 2**, respectively. The iPSC-BMECs on day 8 of differentiation were seeded on the Transwell insert that was pre-coated with collagen IV and fibronectin for a further two days, which helps to purify the BMECs, and then co-cultured with MCs, astrocytes and neurons that were growing on the bottom of the Transwells (**Figure 2A**), respectively. The TEER values were then measured over five consecutive days. Results showed that iPSC derived MCs, astrocytes and neurons could all significantly enhance the TEER values of the iPSC-BMECs (**Figure 2B & 2C**). However, MCs from iPSCs of both CADASIL patients carrying *NOTCH3*^R153C^ and *NOTCH3*^C244Y^ both failed to enhance the barrier function of the iPSC-BMECs (**Figure 2D & 2E**).

**Figure 2.**
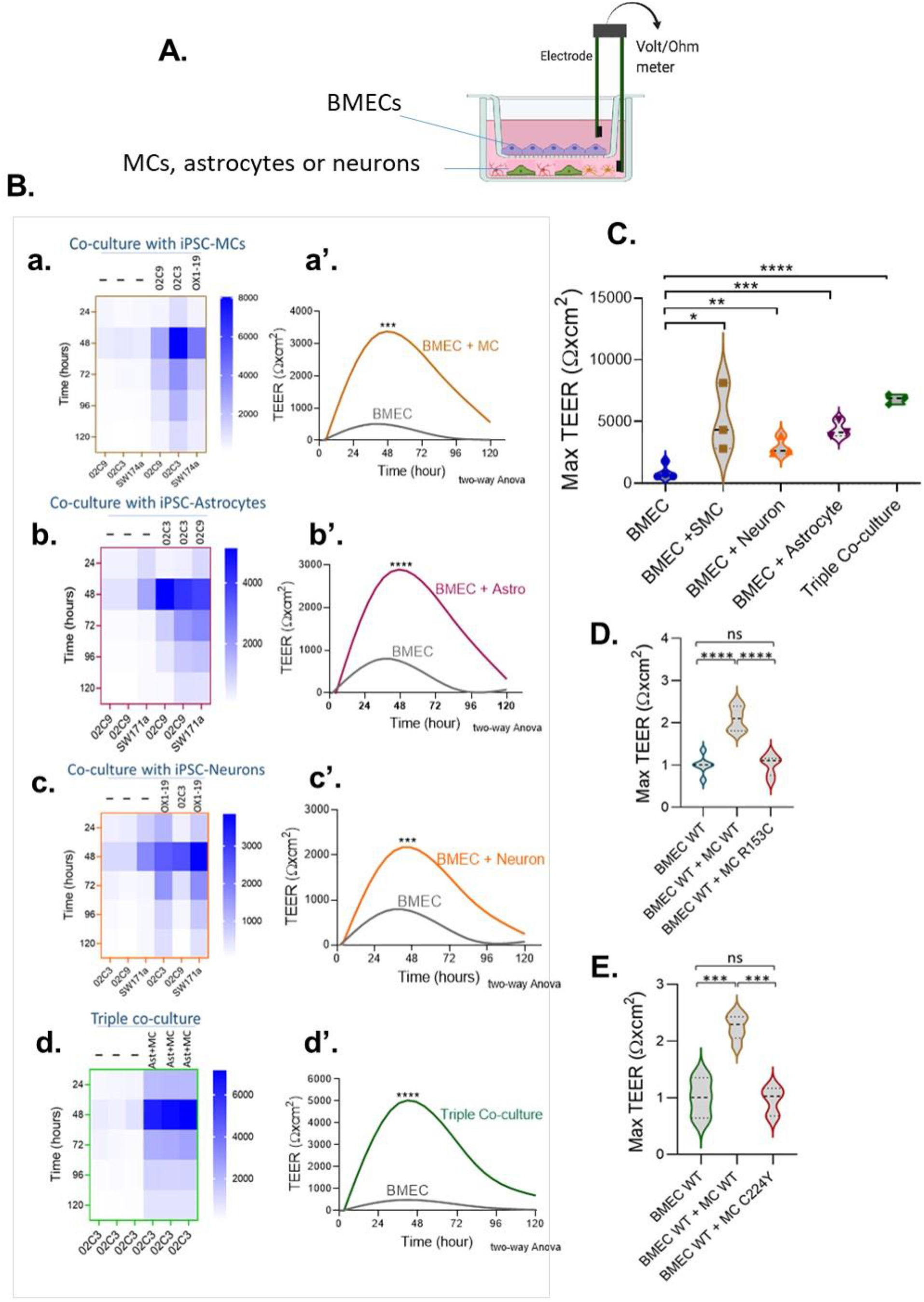
The regulation of BMEC barrier function by iPSC-derived MCs, astrocytes and neurons. **A**. Schematic illustration of the Transwell co-culture NVU system. **B**. Wild-type (WT) iPSC-BMECs were seeded on the insert of Transwell and co-cultured with the WT iPSC derived MCs (**a** and **a’**), astrocytes (**b** and **b’**) and neurons (**c** and **c’**), as well as both astrocytes and neurons (Triple co-coculture, **d** and **d’**) that were grown on the bottom of the Transwell plate. TEER values were measured each day for 5 consecutive days. **Aa.-d**., Heatmaps represent raw TEER values of independent experiments with or without co-culture. A**a’.-d’**., Fit spline curves demonstrate a trend of differences between the BMECs mono-culture and co-culture groups throughout 120 minutes. Two-way ANOVA before post hoc test demonstrated an overall differences between the two groups (***p<0.001, ****p<0.0001). **C**. Comparison of maximum TEER values between groups presented in B. **D** & **E**, BMECs derived from WT iPSCs were co-cultured with MCs from either WT iPSCs or CADASIL *NOTCH3* variants R153C (**D**.) or R224Y (**E**.), and maximum TEER values were compared. Data are presented as mean ± SEM, n=3 in **C, D**, and **E**, and One-way ANOVA with Tukey’s post hoc test, *p<0.05, **p<0.01, ***p<0.001, and ****p<0.0001, n=3. Figure A was created using BioRender.

### 3.2 The barrier function of BMECs derived from CADASIL iPSCs is impaired

Having validated the functionalities of the iPSC-derived neurovascular cells and their interactions, we went on to determine if the barrier function of BMECs derived from iPSCs of CADASIL patients was impaired. Interestingly, we consistently observed low TEER values of iPSC-BMECs from both CADASIL patients as compared with the controls (**Figure 3**). This finding is supported by the immunofluorescent staining where the tight junction proteins (Claudin 5 and Occludin) in the iPSC-BMECs from the two CADASIL patients were significantly disorganised compared to the control iPSC-BMECs (**Figure 4**), although ZO-1 and Glut-1 did not show significant mis-localisation (**Figure 4**).

**Figure 3.**
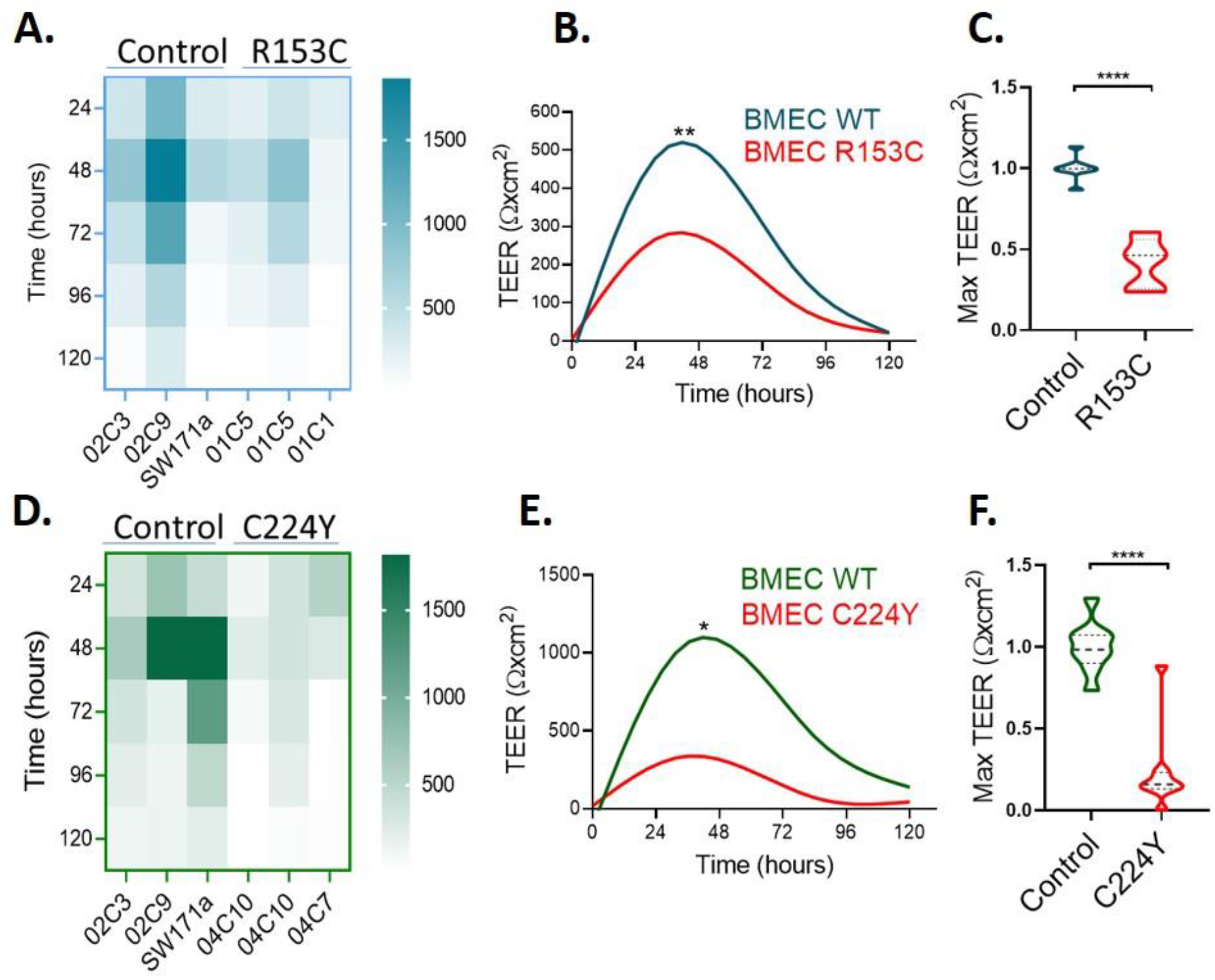
TEER measurement on BMECs generated from iPSCs of CADASIL patients. IPSCs from CADASIL patients carrying *NOTCH3* variants R153C (**A-C**) or C224Y (**D-F**) and healthy control individuals were differentiated into BMECs. TEER values were measured for five consecutive days (120 hours). **A** & **D**, Heatmaps represent raw TEER values of independent experiments for *NOTCH3* (**A**) and *NOTCH3* (**D**). Fit spline curves (**B** & **E**) show a trend of difference between the controls (WT) and mutant groups. Two-way ANOVA before post hoc test demonstrated an overall differences between the two groups (*p<0.05, **p<0.01). **C** & **F**, Maximum TEER values were compared between the control and CADASIL patients carrying *NOTCH3* variant R153C (**C**) and C224Y (**F**). Data are presented as mean ± SEM. Student’s *t*-test, ****p<0.0001, n=3.

**Figure 4.**
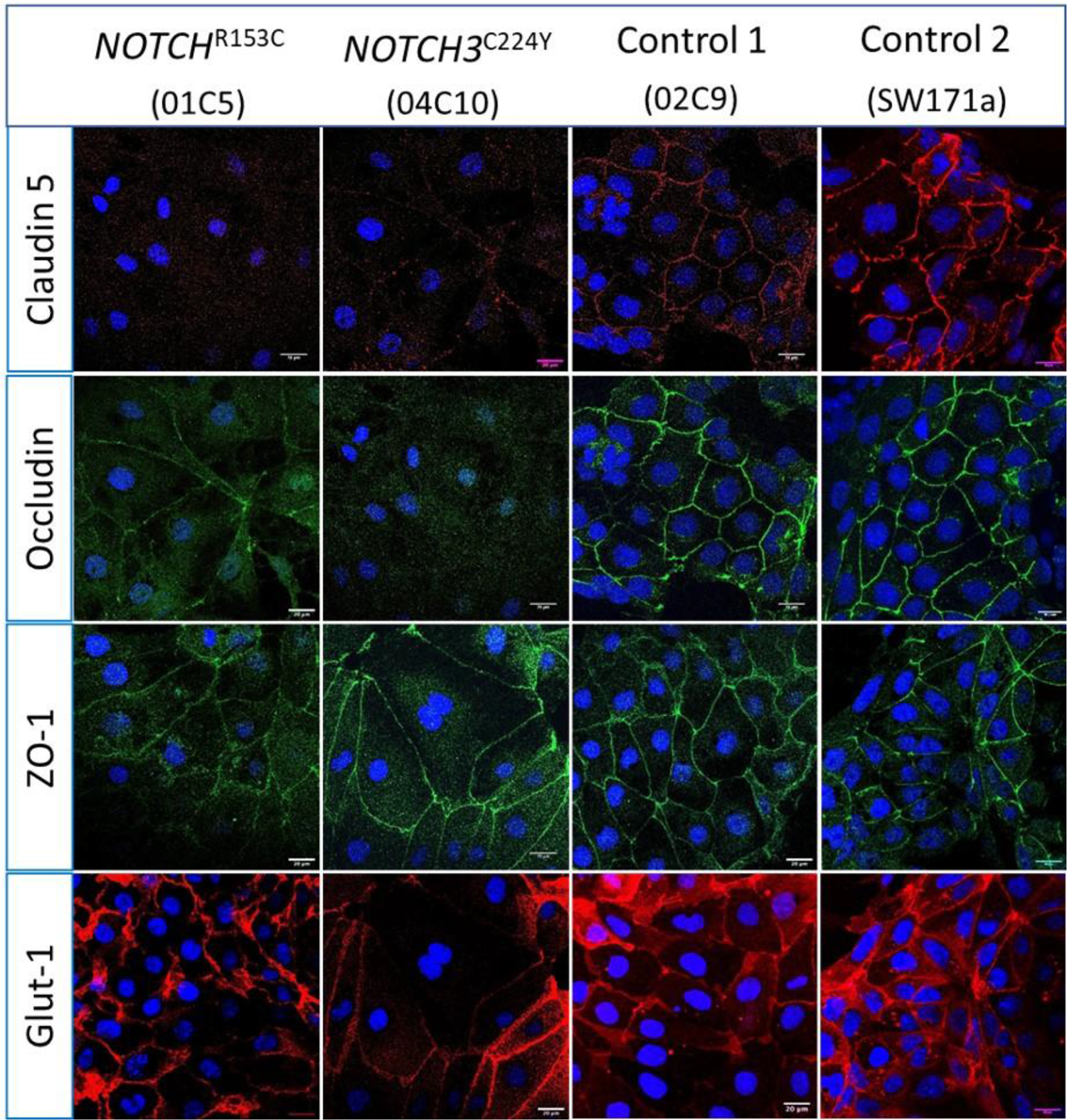
Immunostaining of tight junction proteins and glucose transporter protein in CADASIL iPSC derived BMECs. IPSCs from CADASIL patients carrying *NOTCH3* variants R153C and C224Y, respectively, were differentiated into BMECs and Immunofuorescent stained for tight junction proteins Claudin 5, Occludin and ZO-1, and glucose transporter Glut-1. Images were acquired using confocal microscopy. Scale bar, 20 μm.

Considering the fact that *NOTCH3* is generally expressed in the MCs including VSMCs and pericytes in human adult, it is intriguing to observe an impaired barrier function in BMECs that carry a *NOTCH3* variant. It is possible that the *NOTCH3* expression is higher in BMECs than in the peripheral ECs, which has not been documented previously. We then compared the *NOTCH3* mRNA levels between the iPSC-BMECs and iPSC-ECs. Indeed, the results showed that *NOTCH3* expression in BMECs was about 2-fold higher than that of ECs (**Figure 5A**), while Claudin 5 was highly expressed in BMECs compared to ECs (**Figure 5B**). The higher expression of *NOTCH3* in BMECs than in the peripheral ECs suggests a possible role of NOTCH3 in the brain endothelial cells, which requires further study in the future.

**Figure 5.**
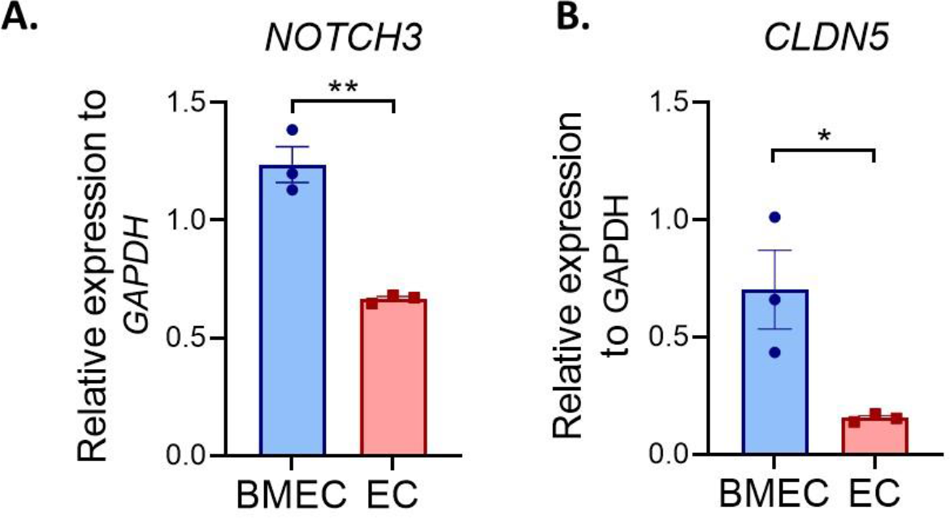
Comparison of *NOTCH3* expression between iPSC derived BMECs and ECs. Wild-type iPSCs differentiated into BMECs and peripheral ECs. The expression of *NOTCH3* and *CLDN5* were determined using qRT-PCR. Data are presented as mean ± SEM. Student’s t-test, *p<0.05, **p<0.001, n=3.

### 3.3 iPSC-derived wild-type MCs, astrocytes and neurons fail to rescue the impaired barrier function of CADASIL iPSC-BMECs

Given the fact that the BBB barrier function of BMECs can be significantly enhanced by MCs, astrocytes and neurons (**Figure 2**), it is interesting to know if the damaged BBB function of the CADASIL BMECs could be rescued by these neurovascular cell types. Thus, the CADASIL iPSC-BMECs were co-cultured with wild type iPSC-derived MCs, astrocytes and neurons, respectively, and TEER values were recorded as described above. Data showed that the TEER of CADASIL iPSC-BMECs could not be rescued by the wild-type iPSC derived MCs (**Figure 6A & 6B**). Astrocytes and neurons seemed to be able to mildly increase the TEER values of the mutant BMECs with statistical significances; however, such function was significantly blunted compared to the co-culture of these cells with the wild-type BMECs (**Figures 6C-F**). These results suggest an intrinsic damage of the BMEC BBB function in CADASIL patients.

**Figure 6.**
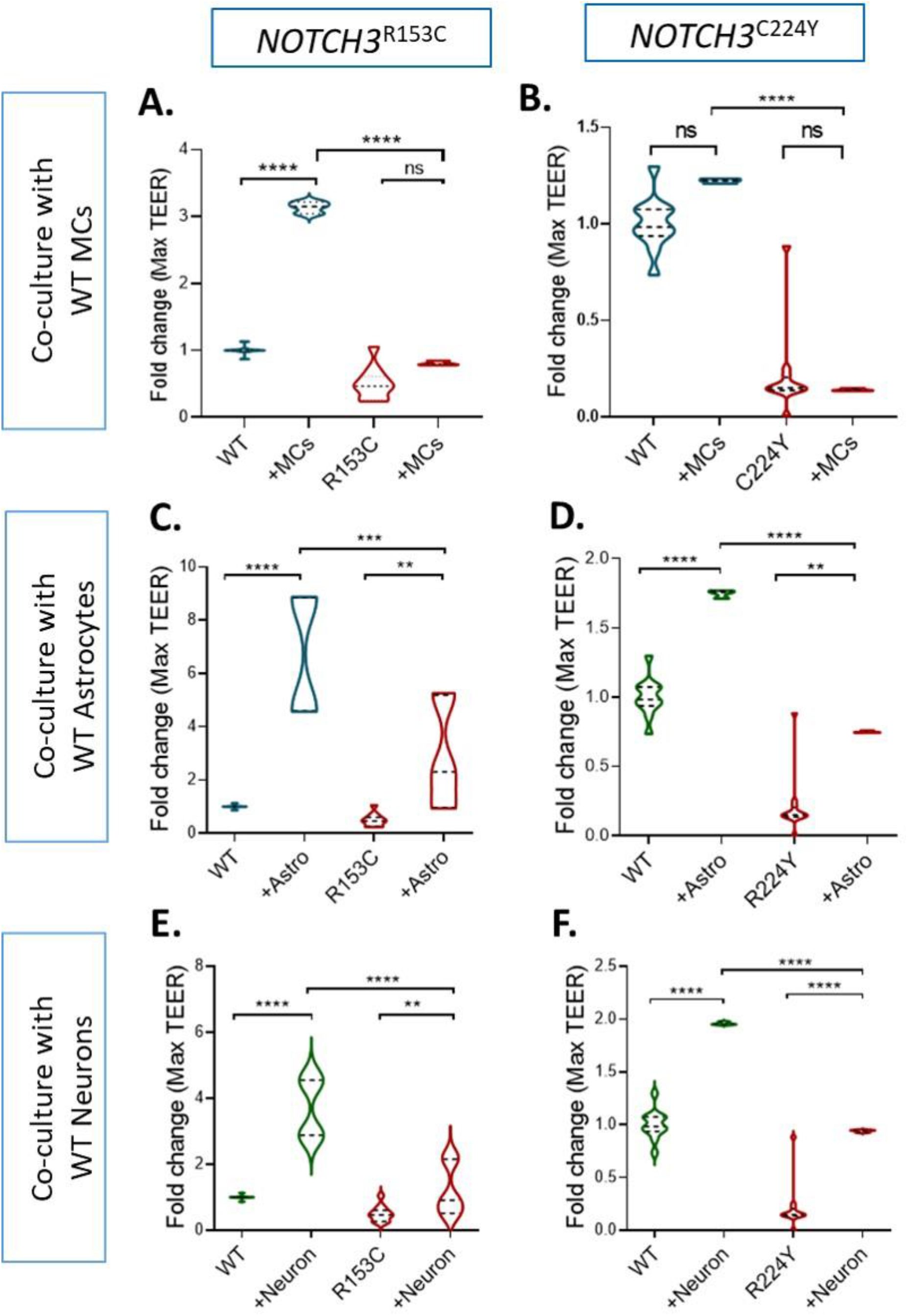
Effect of wild-type iPSC-derived MCs, astrocytes and neurons on the barrier function of CADASIL iPSC-BMECs. **A**. CADASIL or wild-type (WT) control iPSC derived BMECs were seeded on the insert of Transwell and co-cultured with the WT iPSC derived MCs (**A** and **B**), astrocytes (**C** and **D**) and neurons (**E** and **F**) that were grown on the bottom of the Transwell plate. TEER values were measured for 5 consecutive days. Maximum TEER values were presented. IPSC lines from two CADASIL patients (*NOTCH3*R153C and *NOTCH3R244Y)* are presented in the figure. Data are mean ± SEM, n=3. Two-way ANOVA with Tukey’s post hoc test, **p<0.01, ***p<0.001, and ****p<0.0001, n=3.

## 4. DISCUSSION

Using iPSCs from CADASIL patients, we established an *in vitro* NVU model and demonstrated a reduced BBB barrier function in the CADASIL iPSC-BMECs (**Figures 1, 3**), which could not be sufficiently rescued by co-culture with other neurovascular cell types including MCs, astrocytes and neurons that were derived from wild types iPSCs (**Figure 6**). Furthermore, the CADASIL iPSC-MC completely failed to enhance the barrier function of the wild type BMECs (**Figure 6A-B**). CADASIL is a late-onset genetic condition with recurrent ischemic strokes starting at an average age of 45 years. However, some non-specific symptoms like migraine can appear as early as teens with abnormal MRI changes, before the onset of strokes. The impaired neurovascular interactions we demonstrated from the iPSC CADASIL model could at least partially explain the early CADASIL pathologies in the brain. It is likely that the impaired neurovascular interaction could render the brain vulnerable to the subsequent stroke episodes, which accelerates the disease progression to cognitive impairment and dementia.

The unique barrier properties of the brain ECs are not predetermined but induced by the neuronal cues during brain development (Daneman and Prat, 2015, Janzer and Raff, 1987, Alvarez et al., 2013). The complex neurovascular interactions are important for proper BBB function, however, it is challenging to directly measure the contribution of each individual neurovascular cell type to the BBB function in the *in vivo* animal models. In this regard, the iPSC model provides a feasible solution. By co-culturing the iPSC-derived neurovascular cell types in different combinations with the iPSC-BMECs, our iPSC model recapitulated the previous understanding that astrocytes, MCs and neurons could all independently enhance the BBB barrier function of the BMECs and have a synergetic effect (**Figure 2**) (Daneman and Prat, 2015).

During early embryonic development, vascular sprouts interact with the emergent pericytes and radial glial cells in the parenchyma, establishing the initial tight junctions and the immature BBB phenotype (Liebner et al., 2011, Daneman et al., 2009, Daneman et al., 2010b). The development and maturation of the BBB continue during late stages of embryonic development and postnatally when astrocytes becoming a more abundant cell type in the parenchyma. Astrocytes secret a range of factors including TGF-β, glial-derived neurotrophic factor (GDNF), FGF2, angiopoetin 1 (ANG1), sonic hedgehog (Shh), and retinoic acid (RA), all of which induce the formation and strengthen the tight junctions of BMECs (Alvarez et al., 2013). Indeed, our results demonstrated a most effective effect of astrocytes in enhancing the TEER value of BMECs comparing to that of the MCs and neurons (**Figure 2C)**. However, we found that the TEER values of the CADASIL iPSC-BMECs could only be mildly resecured by the wild-type iPSC-astrocytes or iPSC-neurons, which is in great contrast to the significant enhancement effect of the wild-type iPSC-astrocytes and neurons to the wild-type iPSC-BMECs (**Figure 6C-F**). This suggested a likelihood of intrinsic defects of BMECs in CADASIL.

Pericytes are another critical player in the maturation and maintenance of the BBB (Armulik et al., 2011, Brown et al., 2019). However, it was found that pericytes do not necessarily induce the BBB barrier per se but are important in the prevention of the “leaky” properties possibly by inhibiting transcytosis and the expression of leukocyte adhesion molecules (LAMs) on ECs (Armulik et al., 2010, Daneman et al., 2010b, Daneman and Prat, 2015, Armulik et al., 2011). We found that the wild-type iPSC-MCs significantly upregulated the TEER of the wild-type iPSC-BMECs, but the CADASIL iPSC-MCs failed to upregulate the TEER formed by iPSC-BMECs (**Figure 2D & 2E**). Although we have not determined the effect of the CADASIL iPSC-MCs on the transcytosis of BMECs, our previous work found that the CADASIL iPSC-MC is the primary driver to destabilise the microvascular network by a reduced expression of PDGFRβ, decreased secretion of VEGF, and inducing apoptosis of the adjacent ECs (Kelleher et al., 2019). It is plausible that the failure of the CADASIL iPSC-MCs to upregulate the BBB function of wild-type iPSC-BMECs could be at least partially due to a damaging effect of the CADASIL iPSC-MCs on the iPSC-BMECs, and that this cannot be reversed by factors form unaffected iPSC-MCs.

*NOTCH3* is considered to be predominantly expressed in the arterial VSMCs and pericytes (Chabriat et al., 2020). However, we found that the BBB function was significantly reduced in the iPSC-BMECs that carry the CADASIL *NOTCH3* variants (**Figure 3**). The results were confirmed by using iPSC lines from two CADASIL patients on multiple independent experiments. The unique properties of the BMECs that distinguish them from the peripheral ECs led us to explore if NOTCH3 is more important in BMECs than in the peripheral ECs. Using qRT-PCR we indeed found a significantly higher expression of *NOTCH3* in the iPSC-BMECs than in the iPSC-ECs, suggesting a likely unique function of NOTCH3 in BMECs. We also knocked down *NOTCH3* in iPSC-BMECs using siRNA and the pilot data demonstrated a reduced TEER value in the NOTCH3 knockdown iPSC-BMECs (data not shown), which implicates a role of NOTCH3 on the BBB barrier function. Interestingly, a recent publication by Dabertrand et al. (Dabertrand et al., 2021) demonstrated a reduced Kir2.1 activity in cerebral capillary endothelial cells, but not in arteriolar endothelial cells, in a CADASIL mouse model, which was due to compromised PIP2 synthesis in the capillary ECs, suggesting the involvement of BMECs in CADASIL pathology. However, to conclude a definite link between the CADASIL *NOTCH3* variants and BMEC function, more work is required in the future.

In summary, we have demonstrated an impaired neurovascular interaction that contributes to a reduced BBB function in the CADASIL iPSC model. The findings contribute to our current knowledge of the molecular mechanisms underlying CADASIL pathologies and shed light on the understanding of the CNS phenotypes resulting from vascular dysfunction in SVD. Further study is required to refine molecular mechanisms underlying the impaired neurovascular interaction and functional damage of the BBB in CADASIL and identify therapeutic targets. The iPSCs derived from CADASIL patients, albeit an *in vitro* model, is a valuable complement to the *in vivo* animal models and represent a useful human model system for the study of SVD.

## Limitations

Our study has limitations. Firstly, the iPSC control lines were derived from healthy individuals, rather than isogenic controls. Therefore, the observed phenotypes caused by the *NOTCH3* variants may be influenced by variations between individuals. To minimise this effect, we have used iPSC lines from three healthy individuals to compare with two patient iPSC lines in different combinations. It would be ideal to replicate the work using isogenic control lines in the future. Secondly, the protocol of iPSC-BMEC differentiation we adopted (Lippmann et al., 2020, Lippmann et al., 2011, Stebbins et al., 2016), may produce cells that have epithelial properties as argued by a recent publication (Lu et al., 2021). Although we detected CD31 and VE-Cadherin in the iPSC-BMECs which represents endothelial properties, the angiogenic capacity was not as good as the peripheral ECs. We acknowledge the challenge to produce true BMECs that have both endothelial properties and a high BBB function. Effort should be made to further improve the BMEC differentiation protocol in the future.

## 6. AUTHOR CONTRIBUTIONS

WZ: Designed and carried out most of the experiments, data analysis

XZ: Carried out experiments and conducted most of data analysis

XQ: Carried out experiments

SK: Provided iPSC lines, edited manuscript

NH: Supervised project, assisted with study design, edited manuscript

TW: Supervised and designed the project, data interpretation and manuscript writing

## 8. ACKNOWLEDGEMENT

We would like to thank the University of Manchester Bioimaging Core Facility for guidance on confocal microscopy and sample preparation. We thank Professor Pankaj Sharma from Institute of Cardiovascular Research, Royal Holloway, University of London and Imperial College London, and Dr. Helen Murphy and Iris Trender-Gerhard from Manchester Centre for Genomic Medicine (MCGM) for helping recruiting CADASIL patients, which contribute to the establishment of the CADASIL iPSC lines, and thank Dr Adam Dickinson for stablishing the CADASIL iPSC lines.

