## Supplemental figures and tables for "Induced pluripotent stem cell model revealed impaired neurovascular interaction in genetic small vessel disease CADASIL"

### 1. Supplementary Figures and Tables

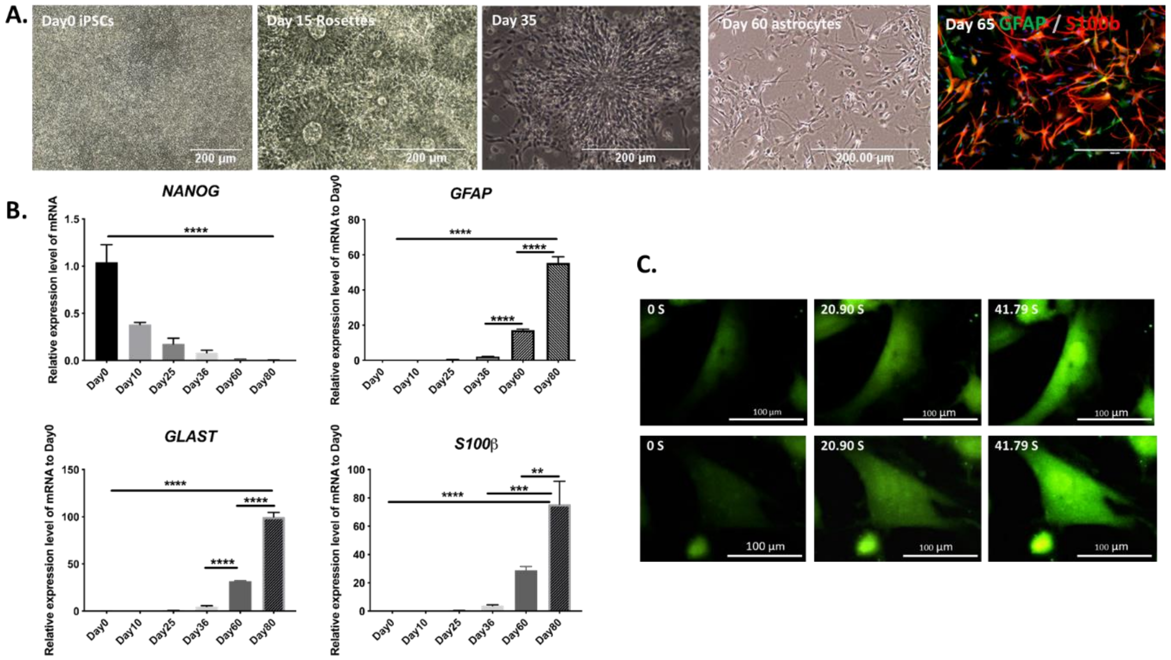


**Suppl. Figure 1**. **Astrocyte differentiation from iPSCs**. **A.** IPSCs were differentiated into astrocytes via a neural progenitor cell stage (neural rosettes shown on day 15 of the differentiation). Images were acquired under light microscope showing morphological changes if iPSC during differentiation up to 60 days. Immunofluorescent staining of the day 65 astrocytes showing the presence of astrocyte markers GFAP and S100b. Scale bar, 200 µm. **B.** qRT-PCR results showing the gradually reduced expression of pluripotent gene *NANOG* and increased expression of astrocyte marker genes *GFAP, GLAST* and *S100b* during the course of astrocyte differentiation from iPSCs up to 60 days. **C.** Calcium imaging showing spontaneous calcium wave in iPSC-derived astrocytes. Scale bar, 100 µm.

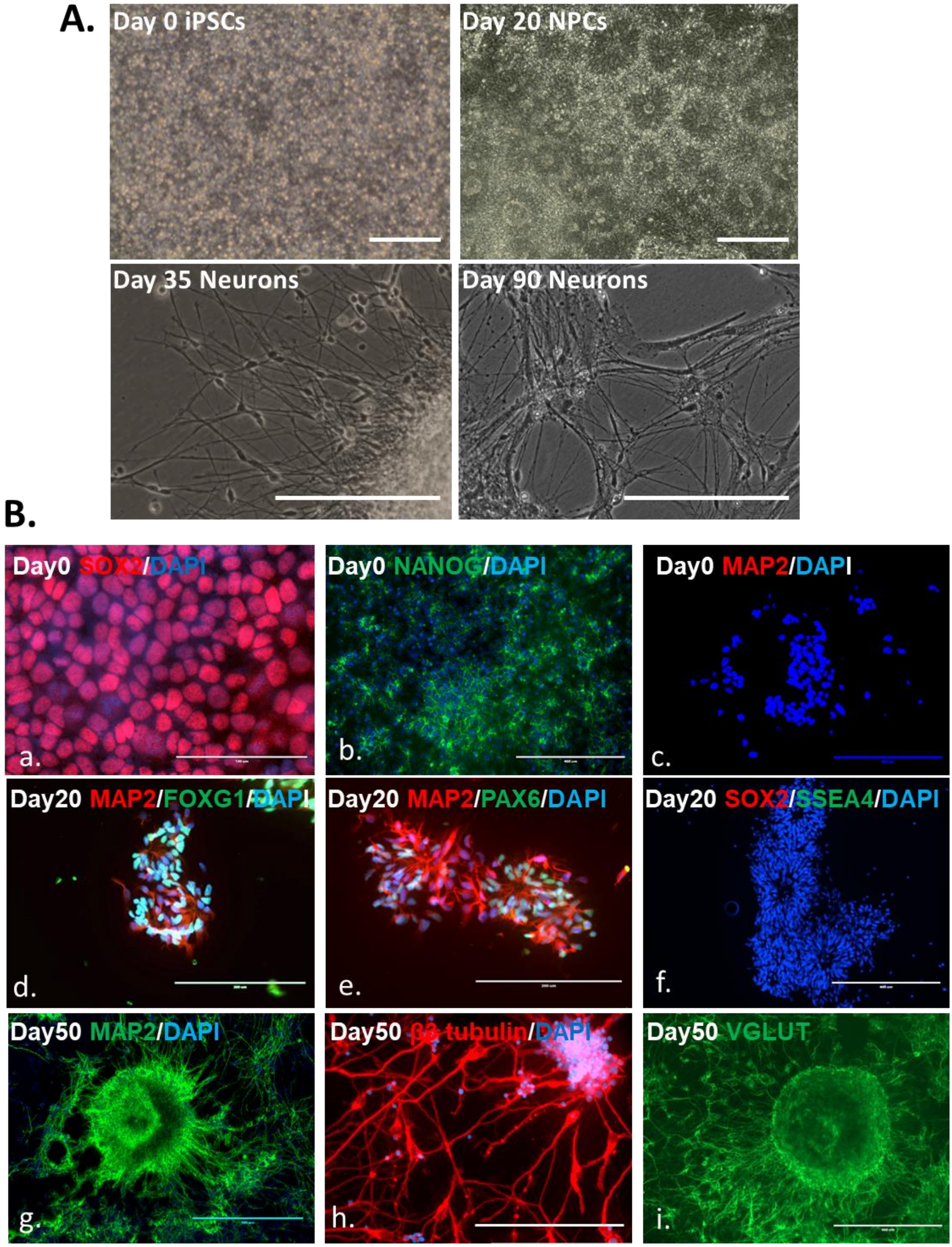


**Suppl. Figure 2**. **Cortical projection neuron differentiation from iPSCs**. **A.** IPSCs were differentiated into cortical projection neurons. Images taken by light microscopy shows morphological changes of the iPSC during the differentiation for up to 90 days. **B**. Immunofluorescent staining showing the presence of marker proteins in cells at different stages of the neuron differentiation. On Day 0, SOX2 and NANOG existed in iPSCs, and neural marker MAP2 was negative; on Day 20, neural progenitor cells displayed MAP, FOXG1 and PAX6, and iPSC marker SOX2 and SSEA4 were negative; on Day 50, neurons displayed neural markers of MAP2, β-tubulin and VGLUT1. Scale bars in A, B.c and B.f-i, 400 μm; in B.d, e and h, 200 μm; in B.a, 100 μm.

#### Suppl. Table 1. Antibodies used for immunofluorescent staining

| Targets | Species | Supplier | Dilution |
|---|---|---|---|
| SOX2 | Rabbit | Abcam ab181557 | 1:500 |
| NANOG | Rabbit | Abcam ab109250 | 1:200 |
| SSEA4 | Mouse | Abcam ab16287 | 1:200 |
| CD31 | Mouse | Bio Techne BBA7 | 1:200 |
| VE-Cadherin | Mouse | Bio Techne MAB9381 | 1:200 |
| Occludin | Mouse | Life Technology 331500 | 1:200 |
| Claudin 5 | Mouse | Life Technology 352500 | 1:100 |
| FOXG1 | Mouse | Abcam ab18259 | 1:500 |
| PAX6 | Mouse | Covance PRB-278B | 1:500 |
| MAP2 | Mouse | Abcam ab92434 | 1:600 |
| β-tubulin | Rabbit | Abcam ab18207 | 1:500 |
| VGLUT1 | Mouse | Life Technology 33150 | 1:200 |
| GFAP | Chicken | Abcam ab4674 | 1:600 |
| S100β | Rabbit | Abcam ab52642 | 1:200 |
| Donkey pAb anti-Rabbit Alexa Fluor® 594 | Donkey | Abcam ab150064 | 1:500 |
| Donkey pAb anti-Mouse Alexa Fluor® 488 | Donkey | Abcam ab150105 | 1:500 |
| Goat pAb anti-Chicken Alexa Fluor® 647 | Donkey | Abcam ab150175 | 1:500 |
| Donkey pAb anti-Goat Alexa Fluor® 488 | Donkey | Abcam ab150029 | 1:500 |

#### Suppl. Table 2. Primer list for qRT-PCR

| Target genes | Forward sequence | Reverse sequence |
|---|---|---|
| *CLDN5* | 5’ GTTCGCCAACATTGTCGTCC 3’ | 5’ GTAGTTCTTCTTGTCGTAGTCGC 3’ |
| *GAPDH* | 5’ CATGTTCGTCATGGGTGTGAACCA 3’ | 5’ ATGGCATGGACTGTGGTCATGAGT 3’ |
| *GFAP* | 5’ GTCCCCCACCTAGTTTGCAG 3’ | 5’ TAGTCGTTGGCTTCGTGCTT 3’ |
| *GLAST* | 5’ ACCCCAAGCATTCTGTGC 3’ | 5’ TTCCGAAATAGAGCCTCGAC 3’ |
| *NANOG* | 5’ TTAATAACCTTGGCTGCCGT 3’ | 5’ GCAGCAAATACGAGACCTCT 3’ |
| *NOTCH3* | 5’ CATCTCCGACCTGATCTGCC 3’ | 5’ GTCTGTAGAGCGGTTTCGGA 3’ |
| *S100b* | 5’ TGTAGACCCTAACCCGGAGG 3’ | 5’ TGCATGGATGAGGAACGCAT 3’ |
